# A comparison of three programming languages for a full-fledged next-generation sequencing tool

**DOI:** 10.1101/558056

**Authors:** Costanza Pascal, Herzeel Charlotte, Verachtert Wilfried

**Affiliations:** imec, ExaScience Life Lab, Kapeldreef 75, 3001 Leuven, Belgium

**Author notes:** Equal contributor (Costanza Pascal, Herzeel Charlotte).

## Abstract

**Background:** elPrep is an established multi-threaded framework for preparing SAM and BAM files in sequencing pipelines. To achieve good performance, its software architecture makes only a single pass through a SAM/BAM file for multiple preparation steps, and keeps sequencing data as much as possible in main memory. Similar to other SAM/BAM tools, management of heap memory is a complex task in elPrep, and it became a serious productivity bottleneck in its original implementation language during recent further development of elPrep. We therefore investigated three alternative programming languages: Go and Java using a concurrent, parallel garbage collector on the one hand, and C++17 using reference counting on the other hand for handling large amounts of heap objects. We reimplemented elPrep in all three languages and benchmarked their runtime performance and memory use.

**Results:** The Go implementation performs best, yielding the best balance between runtime performance and memory use. While the Java benchmarks report a somewhat faster runtime than the Go benchmarks, the memory use of the Java runs is significantly higher. The C++17 benchmarks run significantly slower than both Go and Java, while using somewhat more memory than the Go runs. Our analysis shows that concurrent, parallel garbage collection is better at managing a large heap of objects than reference counting in our case.

**Conclusions:** Based on our benchmark results, we selected Go as our new implementation language for elPrep, and recommend considering Go as a good candidate for developing other bioinformatics tools for processing SAM/BAM data as well.

## Background

The sequence alignment/map format (SAM/BAM) [1] is the de facto standard in the bioinformatics community for storing mapped sequencing data. There exists a large body of work on tools for processing SAM/BAM files for analysis [1, 2, 3, 4, 5, 6, 7, 8, 9, 10, 11, 12, 13, 14, 15]. The SAMtools [1], Picard [2], and GATK [3] software packages developed by the Broad and Sanger institutes are considered to be reference implementations for many operations on SAM/BAM files, examples of which include sorting reads, marking PCR and optical duplicates, recalibrating base quality scores, indel realignment, and various filtering options, which typically precede variant calling. Many alternative software packages [4, 5, 6, 7, 8, 9, 10, 12, 14, 15] focus on optimizing the computations of these operations, either by providing alternative algorithms, or by using parallelization, distribution, or other optimization techniques specific to their implementation language, which is often C, C++, or Java.

We have developed elPrep [8,16], an open-source, multi-threaded framework for processing SAM/BAM files in sequencing pipelines, especially designed for optimizing computational performance. It can be used as a drop-in replacement for many operations implemented by SAMtools, Picard, and GATK, while producing identical results [8,16]. elPrep allows users to specify arbitrary combinations of SAM/BAM operations as a single pipeline in one command line. elPrep’s unique software architecture then ensures that running such a pipeline requires only a single pass through the SAM/BAM file, no matter how many operations are specified. The framework takes care of merging and parallelizing the execution of the operations, which significantly speeds up the overall execution of a pipeline.

In contrast, related work focuses on optimizing individual SAM/BAM operations, but we have shown that our approach of merging operations outperforms this strategy [8]. For example, compared to using GATK4, elPrep executes the 4-step Broad Best Practices pipeline [17] (consisting of sorting, marking PCR and optical duplicates, and base quality score recalibration and application) up to 13x faster on whole-exome data, and up to 7.4x faster on whole-genome data, while utilizing fewer compute resources [8].

All SAM/BAM tools have in common that they need to manipulate large amounts of data, as SAM/BAM files easily take up 10-100GB in compressed form. Some tools implement data structures that spill to disk when reaching a certain threshold on RAM use, but elPrep uses a strategy where data is split upfront into chunks that are processed entirely in memory to avoid repeated file I/O [16]. Our benchmarks show that elPrep’s representation of SAM/BAM data is more efficient than, for example, GATK4, as elPrep uses less memory for loading the same number of reads from a SAM/BAM file in memory [8]. However, since elPrep does not provide data structures that spill to disk, elPrep currently requires a fixed minimum amount of RAM to process a whole-exome or whole-genome file, whereas other tools sometimes allow putting a cap on the RAM use by using disk space instead. Nonetheless, for efficiency, it is recommended to use as much RAM as available [8,18]. This means that, in general, tools for processing SAM/BAM data need to be able to manipulate large amounts of allocated memory.

In most programming languages, there exist more or less similar ways to explicitly or implicitly allocate memory for heap objects which, unlike stack values, are not bound to the lifetimes of function or method invocations. However, programming languages strongly differ in how memory for heap objects is subsequently deallocated. A detailed discussion can be found in “The Garbage Collection Handbook” by Jones, Hosking, and Moss [19]. There are mainly three approaches:

**Manual memory management** Memory has to be explicitly deallocated in the program source code (for example by calling `free` in C [20]).
**Garbage collection** Memory is automatically managed by a separate component of the runtime library called the *garbage collector*. At arbitrary points in time, it traverses the object graph to determine which objects are still directly or indirectly accessible by the running program, and deallocates inaccessible objects. This ensures that object lifetimes do not have to be explicitly modelled, and that pointers can be more freely passed around in a program. Most garbage collector implementations interrupt the running program and only allow it to continue executing after garbage collection – they “stop the world” [19] – and perform object graph traversal using a sequential algorithm. However, advanced implementation techniques, as employed by Java [21] and Go [22], include traversing the object graph *concurrently* with the running program while limiting its interruption as far as possible; and using a multi-threaded *parallel* algorithm that significantly speeds up garbage collection on modern multicore processors.
**Reference counting** Memory is managed by maintaining a reference count with each heap object. When pointers are assigned to each other, these reference counts are increased or decreased to keep track of how many pointers refer to each object. Whenever a reference count drops to zero, the corresponding object can be deallocated.^1^

elPrep was originally, up to version 2.6, implemented in the Common Lisp programming language [23]. Most existing Common Lisp implementations use stop-the-world, sequential garbage collectors. To achieve good performance, it was therefore necessary to explicitly control how often and when the garbage collector would run to avoid needless interruptions of the main program, especially during parallel phases. As a consequence, we also had to avoid unnecessary memory allocations, and reuse already allocated memory as far as possible, to reduce the number of garbage collector runs. However, our more recent attempts to add more functionality to elPrep (like optical duplicate marking, base quality score recalibration, and so on) required allocating additional memory for these new steps, and it became an even more complex task and a serious productivity bottleneck to keep memory allocation and garbage collection in check. We therefore started to look for a different programming language using an alternative memory management approach to continue developing elPrep and still achieve good performance.

Existing literature on comparing programming languages and their implementations for performance typically focus on specific algorithms or kernels in isolation, no matter whether they cover specific domains like bioinformatics [24], economics [25], or numerical computing [26], or are about programming languages in general [27, 28, 29, 30, 31]. Except for one of those articles [31], none of them consider parallel algorithms. Online resources that compare programming language performance also focus on algorithms and kernels in isolation [32]. elPrep’s performance stems both from efficient parallel algorithms for steps like parallel sorting or concurrent duplicate marking, but also from the overall software architecture that organizes these steps into a single-pass, multi-threaded pipeline. Since such software-architectural aspects are not covered by the existing literature, it therefore became necessary to perform the study described in this article.

elPrep is an open-ended software framework that allows for arbitrary combinations of different functional steps in a pipeline, like duplicate marking, sorting reads, replacing read groups, and so on; additionally, elPrep also accommodates functional steps provided by third-party tool writers. This openness makes it difficult to precisely determine the lifetime of allocated objects during a program run. It is known that manual memory management can contribute to extremely low productivity when developing such software frameworks. See for example the IBM San Francisco project, where a transition from C++ with manual memory management to Java with garbage collection led to an estimated 300% productivity increase [33].

Therefore, manual memory management is not a practical candidate for elPrep, and concurrent, parallel garbage collection and reference counting are the only remaining alternatives. By restricting ourselves to mature programming languages where we can expect long-term community support, we identified Java and Go as the only candidates with support for concurrent, parallel garbage collection^2^, and C++17 [36] as the only candidate with support for reference counting (through the `std::shared_ptr` library feature).^3^

The study consisted of reimplementations of elPrep in C++17, Go, and Java, and benchmarking their runtime performance and memory usage. These are full-fledged applications in the sense that they fully support a typical preparation pipeline for variant calling consisting of sorting reads, duplicate marking, and a few other commonly used steps. While these three reimplementations of elPrep only support a limited set of functionality, in each case the software architecture could be completed with additional effort to support all features of elPrep version 2.6 and beyond.

## Results

Running a typical preparation pipeline using elPrep’s software architecture in the three selected programming languages shows that the Go implementation performs best, followed by the Java implementation, and then the C++17 implementation.^4^

To determine this result, we used a five-step preparation pipeline, as defined in our previous article [16], on a whole-exome data set (NA12878). This preparation pipeline consists of the following steps:

- Sorting reads for coordinate order.
- Removing unmapped reads.
- Marking duplicate reads.
- Replacing read groups.
- Reordering and filtering the sequence dictionary.

We ran this pipeline 30 times for each implementation, and recorded the elapsed wall-clock time and maximum memory use for each run using the Unix `time` command. We then determined the standard deviation and confidence intervals for each set of runs [38].

C++17 and Java allow for fine-grained tuning of their memory management, leading to four variations each. For the final ranking in this section, we have chosen the best result from each set of variations, one for C++17 and one for Java. The other results are presented in the Discussion section below. The Go benchmarks were executed with default settings.

The benchmark results for the runtime performance of the three selected implementations are shown in Figure 1. Go needs on average 7 mins 56.152 secs with a standard deviation of 8.571 secs; Java needs on average 6 mins 54.546 secs with a standard deviation of 5.376 secs; and C++17 needs on average 10 mins 23.603 secs with a standard deviation of 22.914 secs. The confidence intervals for Go and Java are very tight, with a slightly looser confidence interval for C++17.

**Figure 1:**
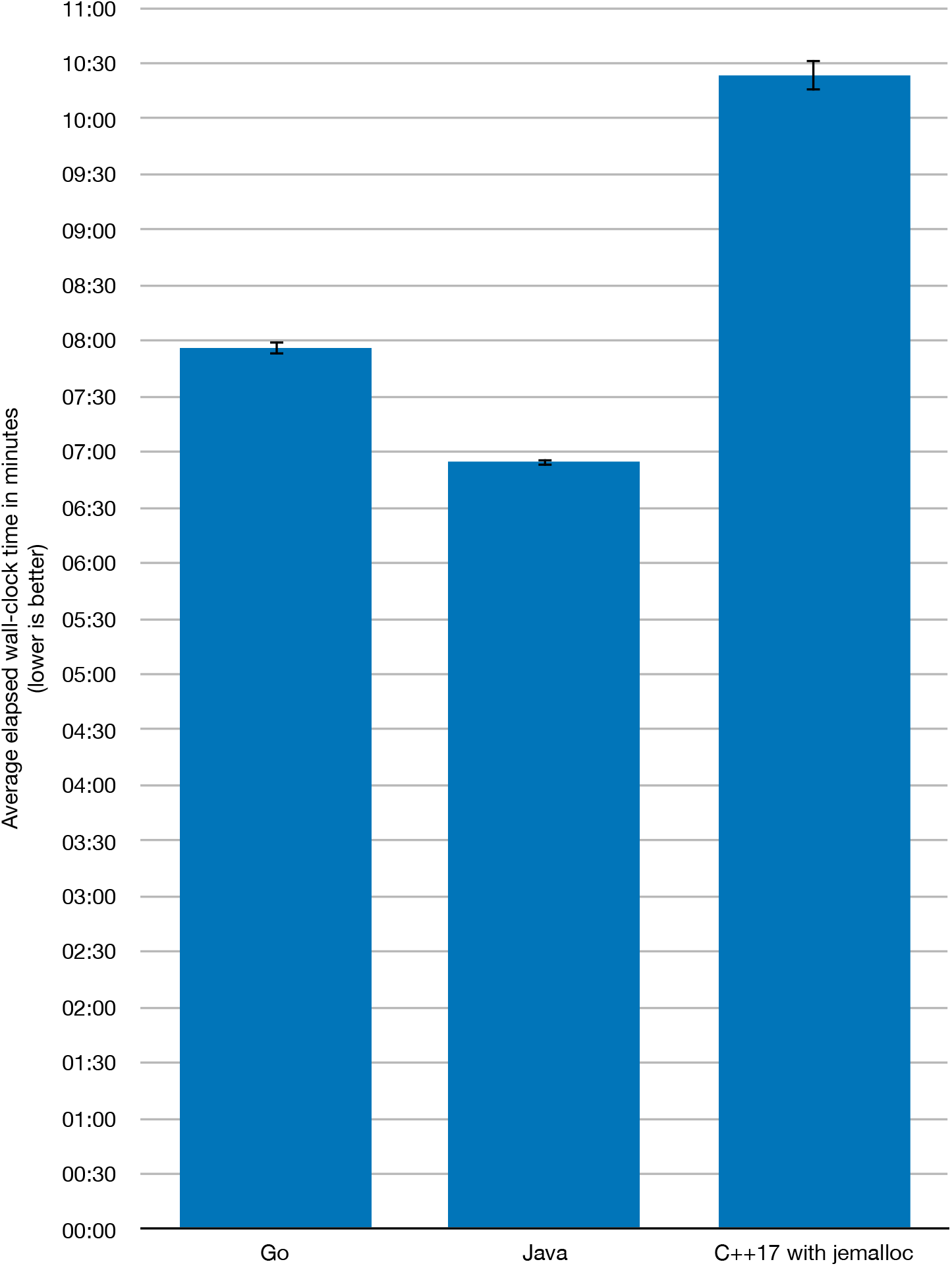
Runtime performance. Average elapsed wall-clock times in minutes for the best Go, Java, and C++17 implementations, with confidence intervals.

The benchmark results for the maximum memory use are shown in Figure 2. Go needs on average ca. 221.73 GB GB with a standard deviation of ca. 6.15 GB; Java needs on average ca. 335.46 GB with a standard deviation of ca. 0.13 GB; and C++17 needs on average ca. 255.48 GB with a standard deviation of ca. 2.93 GB. Confidence intervals are very tight.

**Figure 2:**
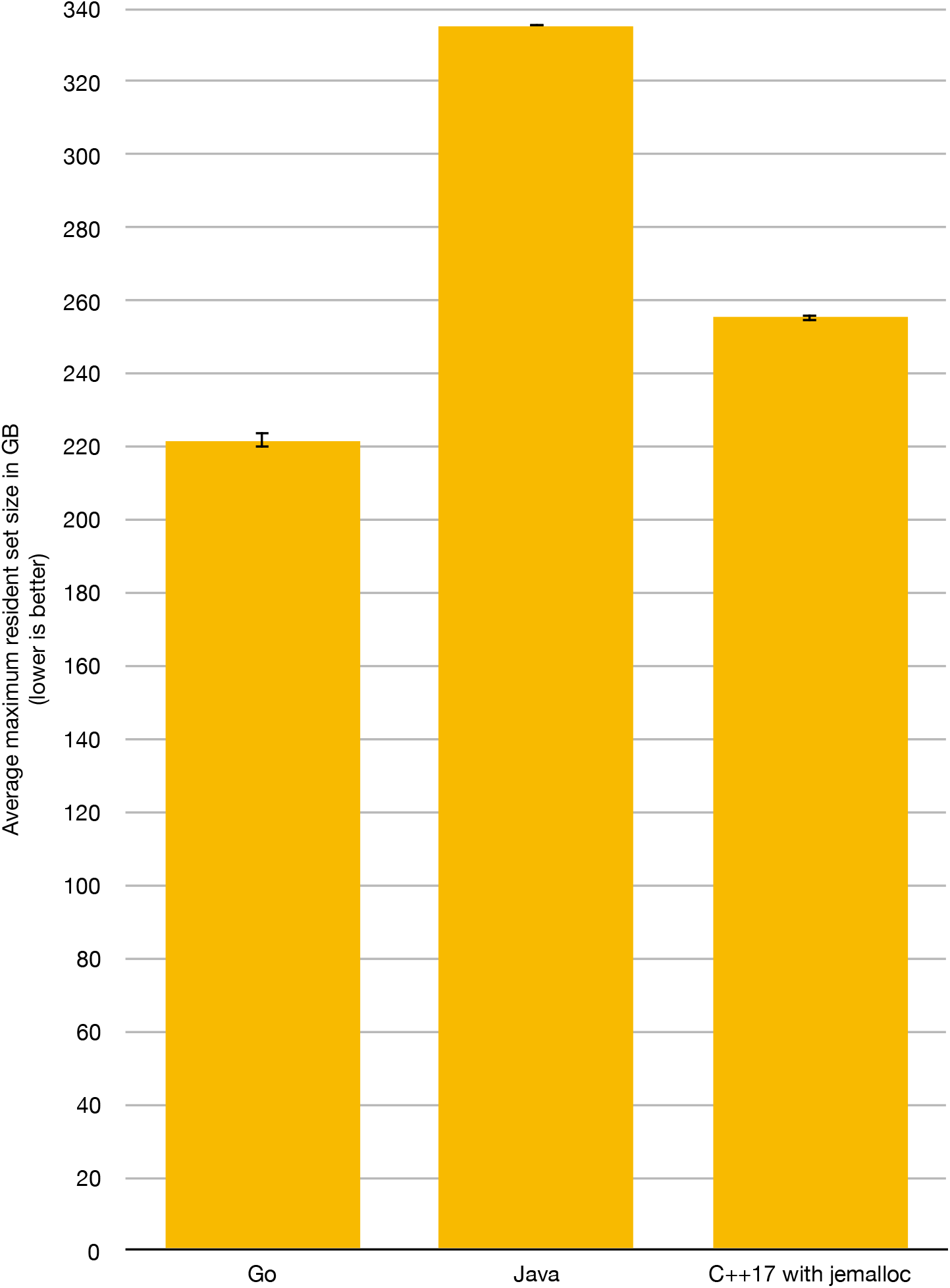
Maximum memory use. Average maximum memory use in GB for the best Go, Java, and C++17 implementations, with confidence intervals

The goal of elPrep is to simultaneously keep both the runtime and the memory use low. To determine the final ranking, we therefore multiply the average elapsed wall-clock time (in hours) with the average maximum memory use (in GB), with lower values (in GBh) being better. This yields the following values (cf. Figure 3):

**Figure 3:**
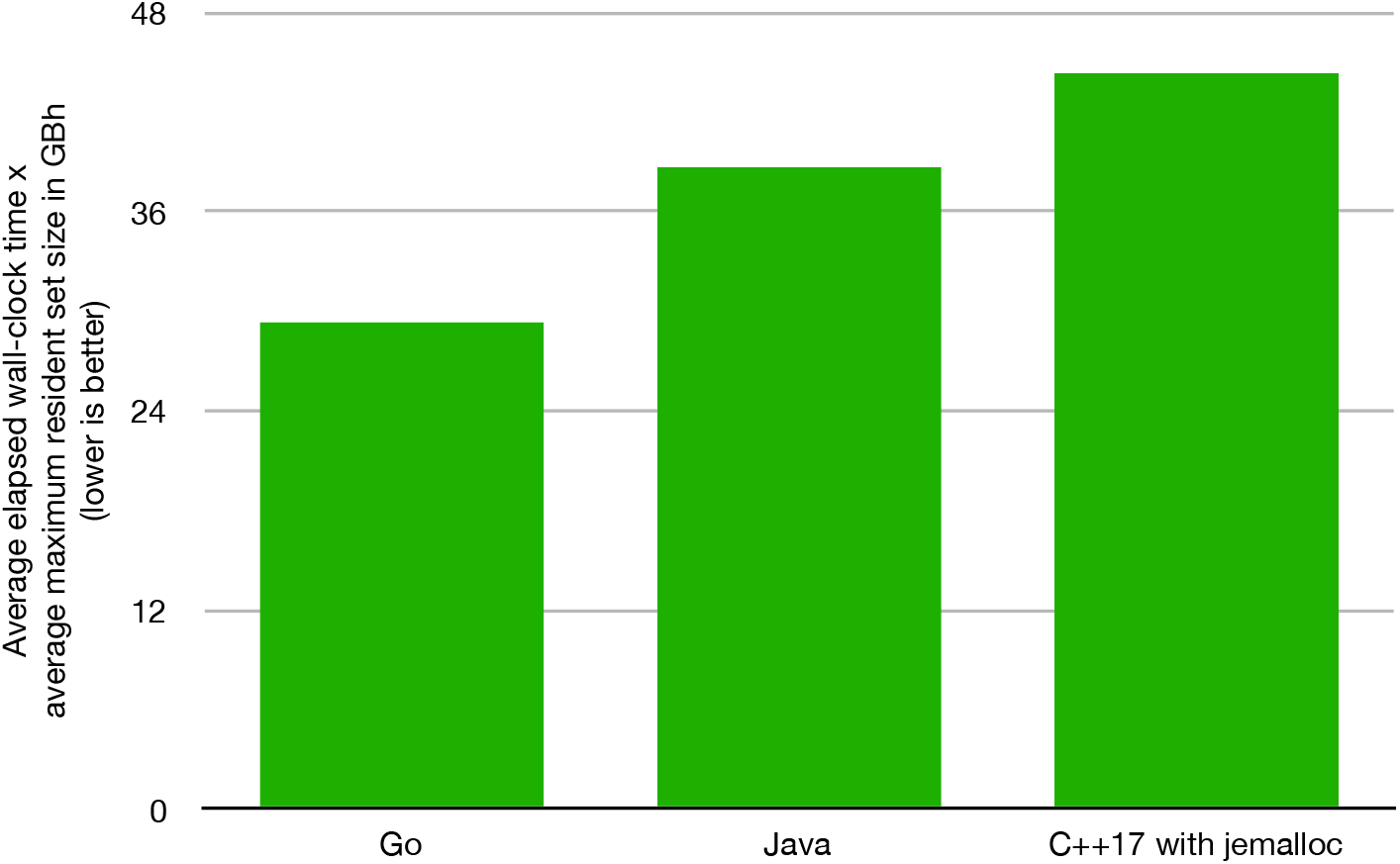
Final ranking of programming languages. Average elapsed wall-clock times multiplied by average maximum memory use in GBh.

- 29.33 GBh for Go
- 38.63 GBh for Java
- 44.26 GBh for C++17

This appropriately reflects the results of the benchmarks: While the Java benchmarks report a somewhat faster runtime than the Go benchmarks, the memory use of the Java runs is significantly higher, leading to a higher GBh value than for the Go runs. The C++17 runs are significantly slower than both Go and Java, explaining the highest reported GBh value. We therefore consider Go to be the best choice, yielding the best balance between runtime performance and memory use, followed by Java and then C++17.

## Discussion

### Memory management issues in elPrep in more detail

The most common use case for elPrep is that it performs sorting of reads and duplicate marking, among other steps [17]. Such a pipeline executes in two phases: In the first phase, elPrep reads a BAM input file, parses the read entries into objects, and performs duplicate marking and some filtering steps on the fly. Once all reads are stored as heap objects in RAM, they are sorted using a parallel sorting algorithm. Finally, in the second phase, the modified reads are converted back into entries for a BAM output file and written back. elPrep splits the processing of reads into these two phases because writing the reads back to an output file can only commence once duplicates are fully known and reads are fully sorted in RAM.

Phase 1 allocates a lot of intermediate data structures for parsing the read representations from BAM files into heap objects. These temporary objects become obsolete after phase 1. The different memory management approaches outlined in the Background section above deal with these temporary objects in different ways.

A garbage collector needs to spend time to classify these obsolete temporary objects as inaccessible and deallocate them. A stop-the-world, sequential garbage collector creates a significant pause in which the main program cannot make progress. This was the case with the previous elPrep versions (up to version 2.6), which is why we provided an option to users to disable garbage collection altogether in those versions [39]. In contrast, a concurrent, parallel garbage collector can perform its job concurrently with phase 2, which can therefore commence immediately.

With reference counting, temporary objects are recognized as obsolete due to their reference counts dropping to zero. Deallocation of these objects leads to transitive deallocations of other objects because of their reference counts transitively dropping to zero. Since this is an inherently sequential process, this leads to a similar significant pause as with a stop-the-world garbage collector.

### C++17 performance in more detail

C and C++ typically perform much better than other programming languages in most benchmarks that focus on isolated algorithms or kernels [24, 25, 26, 28, 29, 30]. Since our C++17 implementation of elPrep uses reference counting, this performance gap may be explained by the deallocation pause caused by reference counting, as described in the previous subsection.

To verify this theory, we timed each phase and the deallocation pause in the C++17 implementation of elPrep separately, and repeated the benchmark another 30 times to determine the timings, standard deviations, and confidence intervals. The results are shown in Figure 4. The first phase needs on average 4 mins 26.657 secs, with a standard deviation of 6.648 secs; the deallocation pause needs on average 2 mins 18.633 secs, with a standard deviation of 4.77 secs; and the second phase needs on average 3 mins 33.832 secs, with a standard deviation of 17.376 secs.

**Figure 4:**
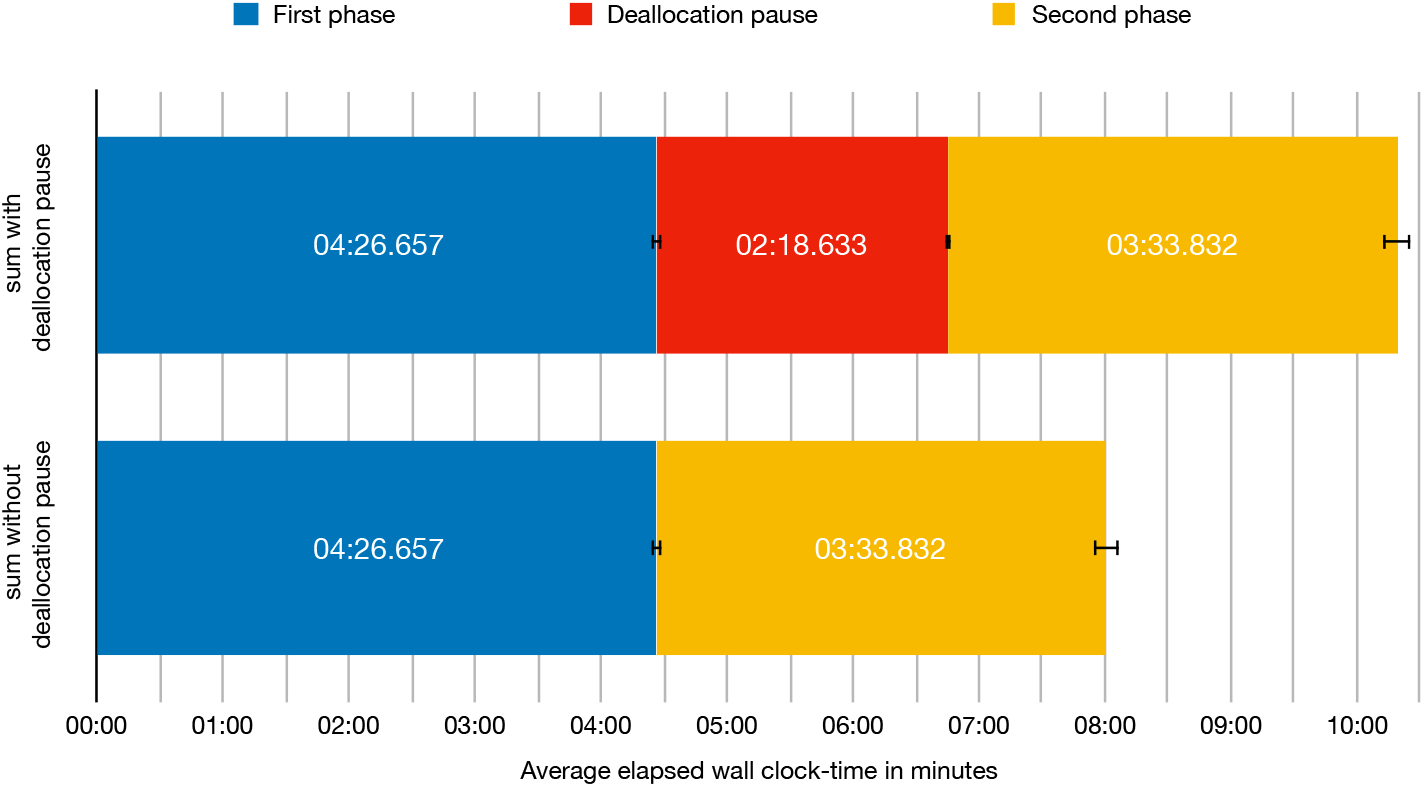
Runtimes of phases in the C++17 implementation. Average elapsed wall-clock times in minutes for the two main phases of an elPrep pipeline in the C++17 implementation, and the deallocation pause in between phase 1 and 2 caused by the reference counting mechanism, with confidence intervals. The second row depicts the same averages as in the first now, but without the deallocation pause. The sum of the two phases in the second row is very close to the Go runtimes shown in Figure 1.

The average total sum of the 30 C++17 runtimes is 10 mins 19.122 secs with a standard deviation of 22.782 secs. If we substract the timings of the deallocation pause from the average total runtime, we get 8 mins 0.489 secs with a standard deviation of 20.605 secs. This is indeed very close to the Go benchmarks which, as reported above, need on average 7 mins 56.152 secs. We therefore conclude that the performance gap between the C++17 version and the Go and Java versions can indeed be explained by the deallocation pause caused by the reference counting mechanism in C++17.

C++ provides many features for more explicit memory management than is possible with reference counting. For example, it provides *allocators* [36] to decouple memory management from handling of objects in containers. In principle, this may make it possible to use such an allocator to allocate temporary objects that are known to become obsolete during the deallocation pause described above. Such an allocator could then be freed instantly, removing the described pause from the runtime. However, this approach would require a very detailed, error-prone analysis which objects must and must not be managed by such an allocator, and would not translate well to other kinds of pipelines beyond this particular use case. Since elPrep’s focus is on being an openended software framework, this approach is therefore not practical.

### Tuning of memory management in C++17

The performance of parallel C/C++ programs often suffers from the low-level memory allocator provided by the C/C++ standard libraries. This can be mitigated by linking a high-level memory allocator into a program that reduces synchronization, false sharing, and memory consumption, among other things [40]. In our study, we have benchmarked the C++17 implementation using the default unmodified memory allocator, the tbbmalloc allocator from Intel Threading Building Blocks [41], the tcmalloc allocator from gperftools [42], and the jemalloc allocator [43]. The measurements are shown in Table 1. According to the listed GBh values, jemalloc performs best.

**Table 1:** Performance results for the different memory allocators used in the C++17 benchmarks: (1) default allocator, (2) tbbmalloc, (3) tcmalloc, (4) jemalloc.

|  | average runtime | average memory | product |
| --- | --- | --- | --- |
| (1) | 16 mins 57.467 secs | 233.63 GB | 66.03 GBh |
| (2) | 16 mins 26.450 secs | 233.51 GB | 63.96 GBh |
| (3) | 11 mins 24.809 secs | 246.78 GB | 46.94 GBh |
| (4) | 10 mins 23.603 secs | 255.48 GB | 44.26 GBh |

### Tuning of memory management in Java

Java provides a number of tuning options for its memory management [44]. Since our Java implementation of elPrep suffers from a significantly higher average maximum memory use than the C++17 and Go implementations, we have investigated two of these options in more detail:

- The string deduplication option identifies strings with the same contents during garbage collection, and subsequently removes the redundancy by letting these strings share the same underlying character arrays. Since a significant portion of read data in SAM/BAM files is represented by strings, it seemed potentially beneficial to use this option.
- The minimum and maximum allowed percentage of free heap space after garbage collection can be configured using the “MinFreeHeap” and “MaxFreeHeap” options to minimze the heap size.

We ran the Java benchmark 30 times each with the following cofigurations: with the default options; with just the string deduplication option; with just the free-heap options; and with both the string deduplication and the free-heap options. For the free-heap options, we followed the recommendation of the Java documentation to reduce the heap size as far as possible without causing too much performance regression. The measurements are shown in Table 2: The free-heap options show no observable impact on the run-time performance or the memory use, and the string deduplication option increases the average elapsed wall-clock time with a minor additional increase in memory use. According to the listed GBh values, Java with default options performs best.

**Table 2:**
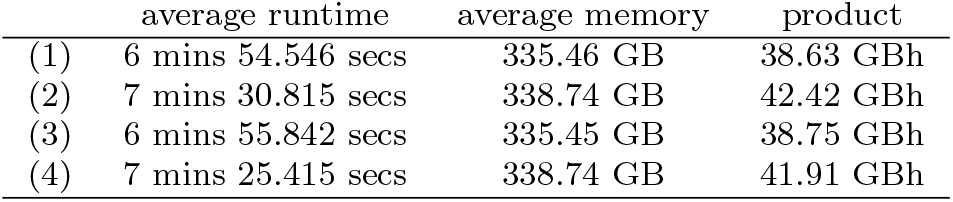
Performance results for the different memory management options used in the Java benchmarks: (1) default options, (2) with string deduplication, (3) with heap-free options, (4) with string deduplication and heap-free options.

## Conclusions

Due to the concurrency and parallelism of Go’s and Java’s garbage collectors, the elPrep reimplementations in these programming languages perform significantly faster than the C++17 implementations which relies on reference counting. Since the Go implementation uses significantly less heap memory than the Java implementation, we therefore decided to base the official elPrep implementation since version 3.0 on Go.

Based on our positive experiences, we recommend authors of other bionformatics tools for processing SAM/BAM data, and potentially also other sequencing data formats, to also consider Go as an implementation language. Previous bioinformatics tools that are implemented in Go include bíogo [45], Fastcov [46], SeqKit [47], and Vcfanno [48], among others.

## Methods

Existing literature on comparing programming languages for performance strives to replicate algorithm or kernel implementations as close to each other as possible across different programming languages, to ensure fair comparisons of the underlying compiler and runtime implementations. We focused on taking advantage of the respective strengths of the different programming languages and their libraries instead. Eventually, a reimplementation of elPrep would have to do this anyway to achieve optimal performance, so this approach results in a more appropriate assessment for our purpose. For example, in C++17 we have used Intel’s Threading Building Blocks as an advanced library for parallel programming, and benchmarked different memory allocators optimized for multi-threaded programs; in Go, we have relied on its concurrency support through *goroutines* and channels for communicating between them; and in Java, we have based elPrep on its framework to support functional-style operations on streams of elements in the package `java.util.Stream` introduced in Java 8.

The benchmarks have all been performed on a Supermicro SuperServer 1029U-TR4T node with two Intel Xeon Gold 6126 processors consisting of 12 processor cores each, clocked at 2.6 GHz, with 384 GB RAM. The operating system used for the benchmarks is the CentOS 7 distribution of Linux. We have used the following compilers and libraries:

- C++17: GNU g++ version 7.2.1 – Threading Building Blocks 2018 Update 2 – gperftools version 2.6.3 – jemalloc version 5.0.1
- Go: Official Go distribution version 1.9.5
- Java: Java Platform, Standard Edition (JDK) 10

For C++17, we additionally used the Intel Threading Building Blocks, gperftools, and jemalloc libraries. The Go and Java versions do not require additional libraries.

We verified that all implementations produce exactly the same results by using the method described in our previous paper on elPrep [16]. This method consists of the following steps:

1. We verify that the resulting BAM file is properly sorted by coordinate order with `samtools index`.
2. We remove the `PG` tag and alphabetically sort the optional fields in each read with `biobambam`.
3. We sort the BAM file by read name and store it in SAM format with `samtools sort`.
4. Finally, we verify that the contents are identical with the result of the original elPrep version with the Unix `diff` command.

## Availability of data and material

The source code for the different elPrep implementations are available at the following locations:

- Common Lisp: https://github.com/exascience/cl-elprep
- C++17, Java: https://github.com/exascience/elprep-bench
- Go: https://github.com/exascience/elprep/tree/v3.04

The five-step preparation pipeline benchmarked in this paper corresponds to the pipeline implemented in the script `run-wes-gatk.sh`, which is available at https://github.com/exascience/elprep/tree/v3.04/demo together with its input files.

## Competing interests

The authors declare that they have no competing interests.

## Authors’ contributions

PC designed and performed the study, participated in the Common Lisp and Go implementations of elPrep, implemented the C++17 and Java versions of elPrep, and drafted the manuscript. CH designed the elPrep software architecture and the benchmarked preparation pipeline, and participated in the Common Lisp and Go implementations of elPrep. PC, CH and WV contributed to the final manuscript.

## Acknowledgements

The authors are grateful to the imec.icon GAP project members, and especially Western Digital for providing the compute infrastructure for performing the benchmarks described in this paper. The authors also thank Thomas J. Ashby and Tom Haber for indepth discussions about memory management techniques in various programming languages.

## Footnotes

1 Object graphs with cycles cannot be easily reclaimed using reference counting alone. However, such cyclic data structures have not occurred yet in elPrep, which is why we do not discuss this issue further in this paper.

2 Specifically, Java uses concurrent, parallel Garbage-first garbage collection [34], whereas Go uses a more traditional concurrent, parallel mark-and-sweep collector [35].

3 Other mature programming languages with support for reference counting include Objective-C, Swift, and Rust [37]. However, in its algorithm for duplicate marking, elPrep requires an atomic compare-and-swap operation on reference-counted pointers, which does not exist in those languages, but exists in C++17.

4 We have not performed a detailed comparison against the original version of elPrep implemented in Common Lisp, but based on previous performance benchmarks, the Go implementation seems to perform close to the Common Lisp implementation.

